# Chromosome loss rate in cells of different ploidy can be explained by spindle self-organization

**DOI:** 10.1101/204230

**Authors:** Ivan Jelenić, Anna M. Selmecki, Liedewij Laan, Nenad Pavin

**Affiliations:** Department of Physics, Faculty of Science, University of Zagreb, Bijenička cesta 32, 10000 Zagreb, Croatia; Department of Medical Microbiology and Immunology, Creighton University Medical School, Omaha, NE; Department of Bionanoscience, Kavli Institute of NanoScience, Faculty of Applied Sciences, Delft University of Technology, P.O. Box 5046, 2600 GA Delft, the Netherlands

## Abstract

The mitotic spindle segregates chromosomes and minimizes chromosome loss for the specific number of chromosomes present in an organism. In *Saccharomyces cerevisiae*, for example, haploid and diploid cells are part of the sexual life cycle and have a thousand times lower rate of chromosome loss than tetraploid cells. Currently it is unclear what constrains the number of chromosomes that can be segregated with high fidelity in an organism. Here we developed a mathematical model to study if different rates of chromosome loss in cells with different ploidy can arise from changes in (1) spindle dynamics and (2) a maximum duration of mitotic arrest, after which cells enter anaphase. Our model reveals how small increases in spindle assembly time can result in exponential differences in rate of chromosomes loss between cells of increasing ploidy and predicts the maximum duration of mitotic arrest.

## INTRODUCTION

Chromosome segregation is an important, highly conserved cellular function. A complex network of interacting components segregates chromosomes with high precision. However, rare errors in chromosome segregation are observed. The error rate generally increases when the number of sets of chromosomes (ploidy) increases within the cell (Comai, 2005). Experimental studies suggest that the spindle is optimized to minimize chromosome loss for the number of chromosomes present in that specific organism (Nannas et al., 2014; Schulman and Bloom, 1993; Storchova et al., 2006). For example, the normal sexual life cycle of the budding yeast *Saccharomyces cerevisiae* includes haploid (*c*= 16 chromosomes) and diploid cells (*c*= 32 chromosomes). In this organism the effect of ploidy on the rate of chromosome loss is very pronounced: haploid and diploid cells have chromosome loss rates around 10^−6^ chromosomes per cell per cell division, whereas tetraploid cells have a loss rate around 10^−3^ (Mayer and Aguilera, 1990; Storchova et al., 2006). Moreover, when ploidy levels are changed in laboratory yeast strains, the ploidy levels tend to return to the initial wild-type level in experimental evolution studies (Gerstein et al., 2006; Selmecki et al., 2015). In the wild, however the number of chromosomes per cell varies substantially between species, suggesting that cells can be optimized for different numbers of chromosomes (Otto and Whitton, 2000).

Chromosome segregation is driven by the mitotic spindle, a self-organized micro machine composed of microtubules and associated proteins. During spindle assembly, spindle poles nucleate microtubules, which grow in random directions or in a direction parallel with the central spindle (O’Toole et al., 1997; Winey et al., 1995). A microtubule that comes into the proximity of a kinetochore (KC), a protein complex at the sister chromatids, can attach to the KC and thus establish a link between chromatids and spindle poles (Akiyoshi et al., 2010; Gonen et al., 2012; Hill, 1985; Mitchison and Kirschner, 1985; Volkov et al., 2013). Theoretical studies have quantitatively shown that this process can contribute to spindle assembly (Kalinina et al., 2013; Paul et al., 2009; Vasileva et al., 2017; Wollman et al., 2005). Prior to chromosome separation, all connections between chromatids and the spindle pole must be established, and erroneous KC-microtubule attachments must be corrected for which several mechanisms are proposed (Tubman et al., 2017; Zaytsev and Grishchuk, 2015). These connections are monitored by the spindle assembly checkpoint (Li and Murray, 1991). Once KCs are properly attached and chromosomes congress to the metaphase plate (Gardner et al., 2008), the spindle assembly checkpoint is silenced and microtubules separate the sister chromatids. Cells have a limited time before progressing to anaphase, after which they enter anaphase regardless of erroneous connections, which we refer to as the maximum duration of mitotic arrest, and can result in chromosome loss (Rieder et al., 1994; Rudner and Murray, 1996). Even though a mechanistic picture of spindle assembly is emerging, it is an open question how changes in ploidy can result in a dramatic effect on the rates of chromosome loss.

## RESULTS

### Model for chromosome loss

In this report we introduce a model for chromosome loss in cells with different ploidy (For description see STAR Methods). We describe populations of cells in prometaphase, metaphase, and anaphase with either all KCs attached to the spindle, or with at least one unattached KC, which can become a lost chromosome (Figure 1A). Transitions between these populations arise from the dynamics in spindle assembly: chromosome attachment, chromosome detachment, and silencing of the spindle assembly checkpoint (Figure 1B). In addition, we introduce a function describing a maximum duration of mitotic arrest after which cell enter anaphase regardless whether all chromosomes are attached. This function allows for chromosome loss in our model.

**Figure 1.**
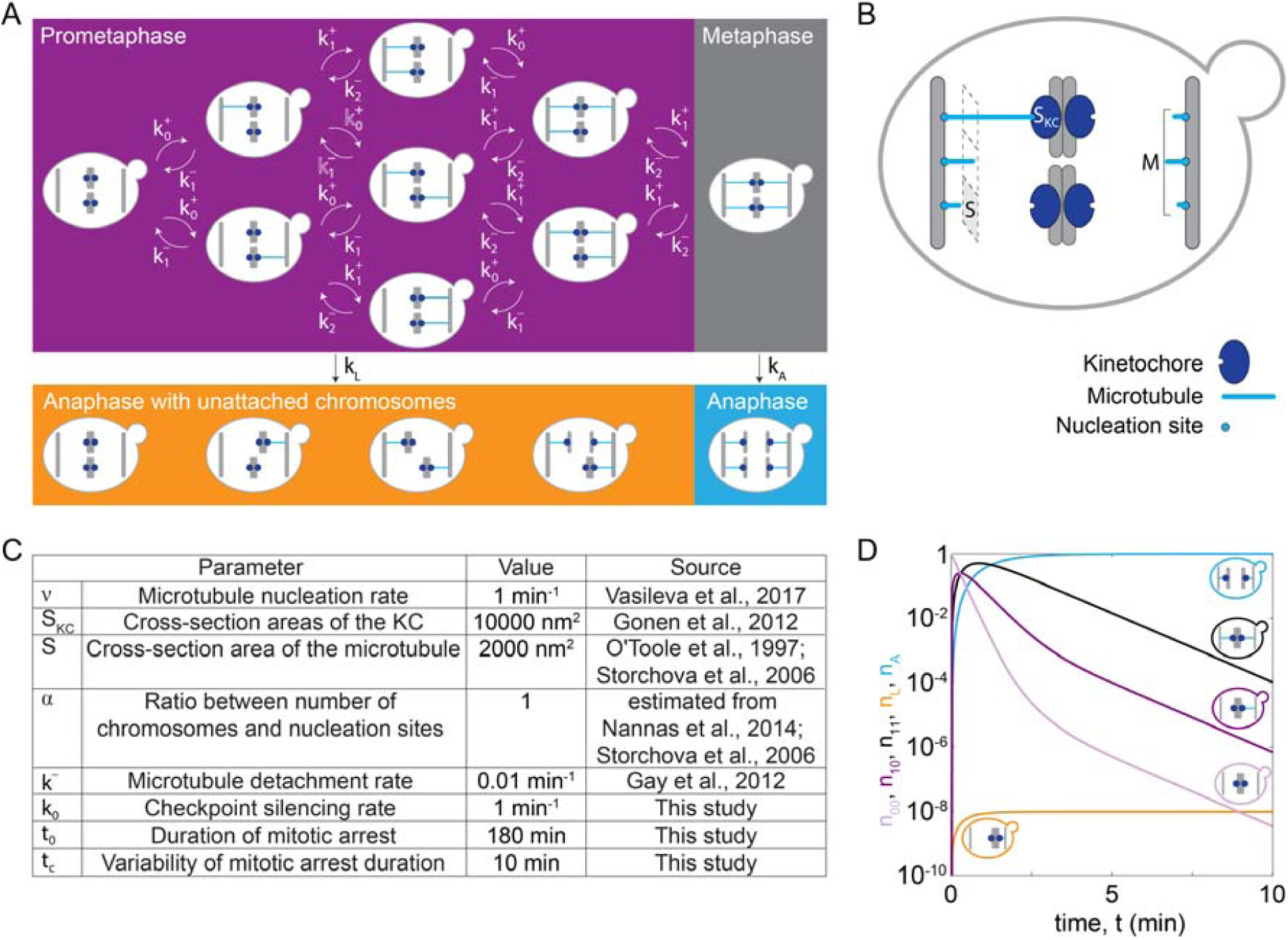
Model for chromosome loss. (A) Scheme of the model: The different boxes indicate cells in prometaphase (purple box), metaphase (grey box) and anaphase (orange and blue box). Arrows denote the rate of transition between different populations. Within a cell, microtubules (blue lines) extend from the spindle pole bodies (grey bars) towards the kinetochores (dark blue circles). (B) Spindle dynamics in an individual cell. A microtubule (light blue) occupies a cross-section area *s*. Microtubules nucleate from *M* nucleation sites at the spindle pole body (grey bar) and extend towards KCs (dark blue) of a cross-section area *s*_KC_. (C) Table 1 lists the parameters used to solve the model. Five parameter values were taken from previous studies (Gay et al., 2012; Gonen et al., 2012; Nannas et al., 2014; O’Toole et al., 1997; Storchova et al., 2006; Vasileva et al., 2017), as indicated. (D) Solution of the model for cells with 1 chromosome (*C*= 1). Fraction of cells in prometaphase with no KCs attached (light purple, *n*_00_), with 1 KC attached (dark purple, *n*_10_), in metaphase (black, *n*_11_), in anaphase with at least one KC unattached (orange, *n*_L_) and in anaphase (blue, *n*_A_), are shown. Each line is accompanied by a cell cartoon depicting the corresponding phase of the cell cycle.

### Time course of chromosome attachment and progression to anaphase

To illustrate how chromosome loss occurs during the transition from prometaphase to anaphase, we numerically solve our model first for cells with only one chromosome, *C*= 1, for parameters given in Figure 1C. We plot the time course for different populations of cells. Initially, cells have no chromosome attached to the spindle, *n*_00_ = 1. In prometaphase, when spindle assembly starts and KCs attach to the spindle, the fraction of cells in this population decreases, while the fraction of cells in other populations increases (compare light and dark purple lines in Figure 1D). After an initial increase, the fraction of cells in prometaphase start decreasing as more KCs attach and cells switch to metaphase (compare purple and black lines in Figure 1D). Finally, cells switch to anaphase. The fractions of cells in anaphase increase and asymptotically approach a limit value because the model does not describe cells leaving anaphase (orange and blue lines in Figure 1D). In this case with only one chromosome, the fraction of cells with a lost KC is very low.

### Model explains dramatic increase in rate of chromosome loss with increase in ploidy

To explore the relevance of our model for haploid, diploid, and tetraploid yeast cells, we further solve our model for *C*= 16, 32, and 64 (Figure 2A). We find that cells with an increasing number of chromosomes spend longer time in prometaphase and metaphase, though the general trend is similar to the case with *C*= 1. However, as time progresses, there is a rapid decrease in the fraction of cells in prometaphase and metaphase which corresponds to reaching the maximum time of mitotic arrest, *t*= *t*_0_. Because populations of cell with more chromosomes have a larger time lag predominantly in prometaphase they also enter anaphase later (Figure 2B). This time lag also results in an increasing fraction of cells in anaphase with at least one lost chromosome because these cells have a greater chance to proceed to anaphase without a completely formed spindle.

**Figure 2.**
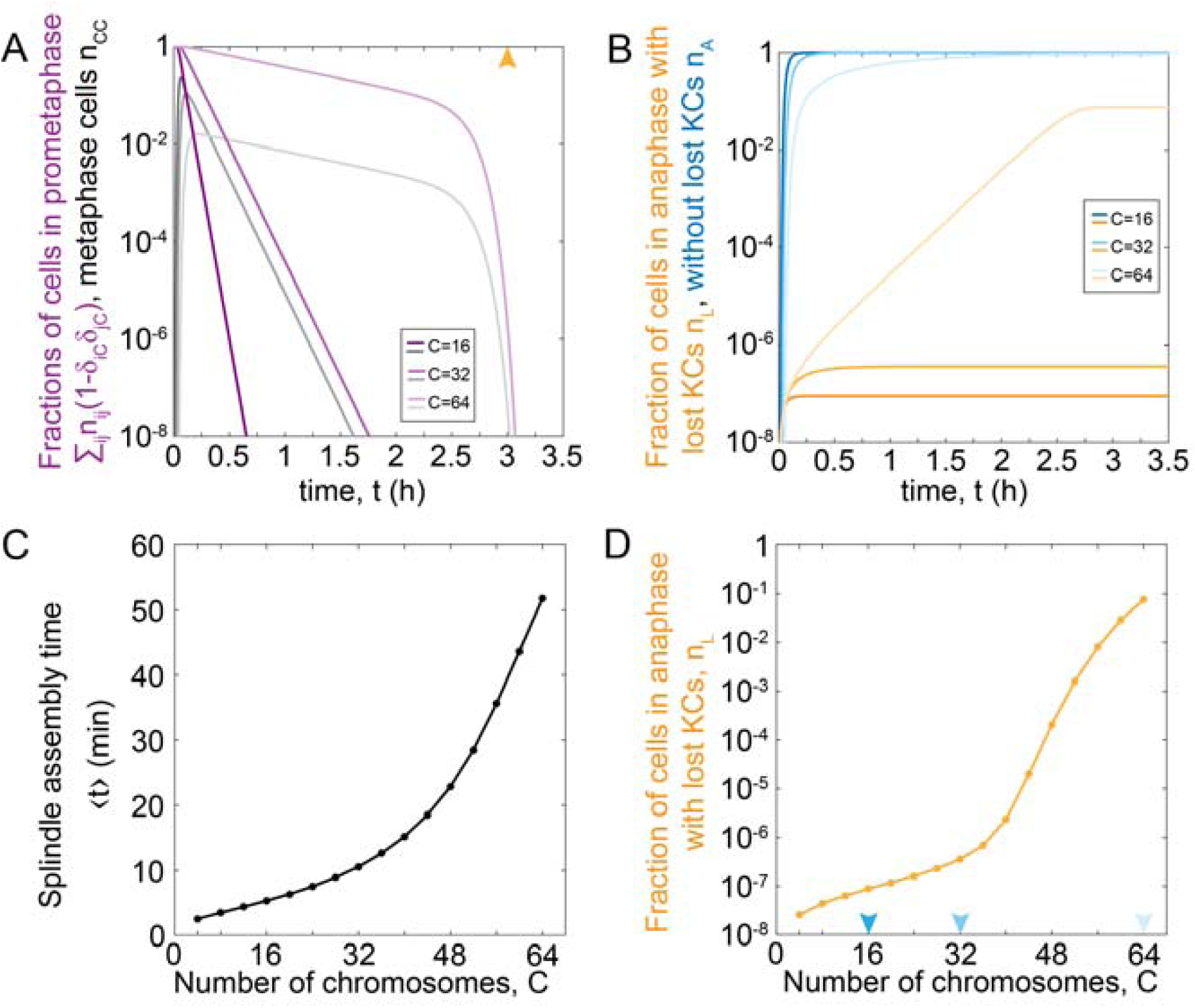
Model predictions for chromosome loss in cells of different ploidy. (A) Fraction of cells in prometaphase (purple) and metaphase (grey) for different numbers of chromosomes. Orange arrowhead denotes value of the duration of mitotic arrest, *t*_0_. (B) Fraction of cells in anaphase with at least one KC unattached (orange) and in anaphase (blue). Three different shades in panels (A) and (B) correspond to different number of chromosomes, *C* = 16,32,64. For color-codes see inset legends. (C) Time of the spindle assembly as function of number of chromosomes. (D) Rate of chromosome loss for cells as function of number of chromosomes. Arrowheads denoted haploid, diploid and tetraploid number of chromosomes. The other parameters are given in Figure 1C.

### Spindle assembly time and the rate of chromosome loss increase with number of chromosomes

To explore which processes in spindle assembly are responsible for significant chromosome loss, we determine the relevance of parameters in our model. As our model describes both spindle formation and transition to anaphase, we separately analyze the contribution of each process. We introduce the average time of both prometaphase and metaphase, which we term the time of spindle assembly (STAR Methods). We find that the time of spindle assembly increases with the number of chromosomes, which we plot on a linear scale (Figure 2C). Cells with more microtubules in the spindle and/or larger KC have shorter time of spindle assembly (Figure S1A, S1B). Next, we explored how ploidy affects chromosome loss. We find that haploid and diploid cells have the same order of magnitude in the fraction of the population with at least one lost chromosome (Figure 2D). Interestingly, the fraction of cells with at least one lost chromosome increases dramatically for cells with higher ploidy, such as tetraploid cells (C=64). This behavior does not qualitatively change by changing parameters describing the duration of mitotic arrest (Figure S2A, S2B). In conclusion, linear changes in spindle assembly time result in exponential differences in chromosome loss rate as soon as prometaphase time approaches the maximum time of mitotic arrest.

## DISCUSSION

Here we introduced a model by which we explored chromosome loss taking into account key aspects of spindle assembly, including microtubule nucleation, KC attachment/detachment, and spindle geometry, together with a maximum time of mitotic arrest. Our theory provides a plausible explanation for experiments in yeast tetraploid cells where there is a thousand-fold increase in the rate of chromosome loss relative to haploid and diploid cells (Mayer and Aguilera, 1990). Our model may also be relevant to mammalian and plant cell types that also experience increased rates of chromosome mis-segregation with increasing ploidy (Hufton and Panopoulou, 2009).

In yeast cells of different ploidy, chromosome loss can occur for many reasons. Our model predicts that longer duration of spindle assembly, which occurs in cells with more chromosomes, increases chromosome loss. This prediction can be verified by measurements of average spindle assembly time in haploid, diploid, and tetraploid yeast cells. Importantly, key parameters of cytoplasmic microtubule dynamics were measured previously for diploid and tetraploid *S. cerevisiae* cells, including the rates of microtubule growth, shrinkage, catastrophe and rescue during G1 and mitosis (Storchova et al., 2006). We hypothesize that change in these parameters may cause an increase in the average spindle assembly time in a population of cells, but experimental validation in yeast is needed.

Laboratory tetraploid yeast cells have an increased rate of chromosome loss. However, a recent experimental evolution study with laboratory yeast found that some tetraploid cell lines could maintain their full chromosome complement (*C*= 64) for >1000 generations (Lu et al., 2016). The evolved, stable tetraploid cells had elevated levels of Sch9 protein, one of the major regulators downstream of TORC1, which is a central controller of cell growth. Interestingly, the evolved stable tetraploid cells also had increased resistance to the microtubule depolymerizing drug benomyl, indicating that increased Sch9 protein activity may at least in part rescue spindle formation defects observed in wild-type tetraploid cells (Lu et al., 2016; Storchova et al., 2006). This is consistent with our model, where chromosome stability in tetraploid cells can be obtained by increasing the rate of spindle assembly.

## STAR+METHODS

Detailed methods are provided in the online version of this paper and include the following:

## SUPPLEMENTAL INFORMATION

Supplemental Information includes two figures and can be found with this article online.

## AUTHOR CONTRIBUTION

N.P., L.L. and A.M.S. conceived and supervised the project. N.P. and L.L. developed the model, I.J. solved the model. All authors wrote the paper.

## ACKNOWLEDGEMENTS

We thank Ivana Šarić for the drawings. N.P. acknowledges support from QuantiXLie Center of Excellence. L.L. gratefully acknowledges support by the Netherlands Organization for Scientific Research (NWO/OCW), as part of the Frontiers of Nanoscience program. This is a pre-print of an article published in Frontiers in Genetics. The final authenticated version is available online at: https://doi.org/10.3389/fgene.2018.00296.

## STAR METHODS

### Model

In our model, we denote the fractions of cells in prometaphase and metaphase by *n*_*ij*_. The fraction of cells in anaphase with at least one unattached KC to the spindle is denoted *n*_L_, and the fraction with all KCs attached is denoted *n*_A_. The indices *i* and *j* denote the number of left and right sister KCs attached to the spindle, respectively, in cells with *C* chromosomes (*i* = 0,…, *C* and *j* = 0,…, *C*). The combination of indices *i*= *j* = *C* describes cells with all KCs attached, which corresponds to metaphase cells. All the other combinations of indices describe cells with at least one unattached KC, which correspond to prometaphase cells. As time, *t*, progresses, cells switch between fractions when (i) KCs attach to or detach from the spindle, or (ii) cells enter anaphase (Figure 1A). We describe these processes by a system of rate equations:

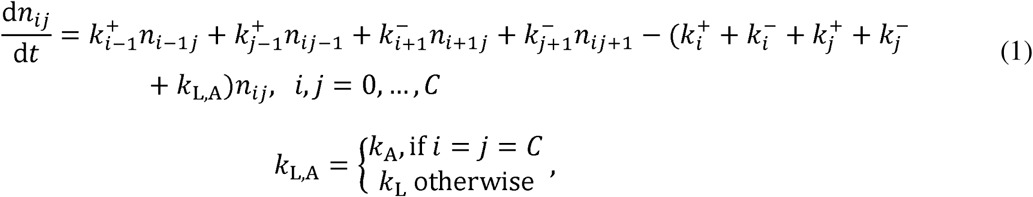

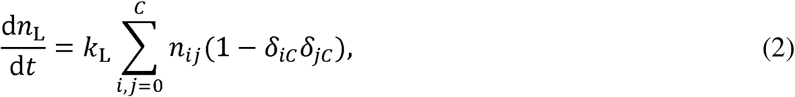

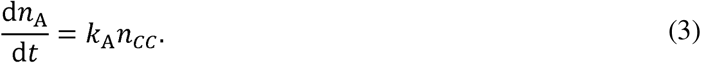

Here, 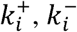, *k*_A_, and *k*_L_ denote the rate of KC attachment, rate of KC detachment, rates of cells entering anaphase with and without all KC attached, respectively, and *δ* denotes the Dirac delta function. The fractions are normalized by the total number of cells in the population, so that 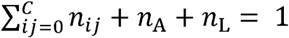. We also introduce the average time of both prometaphase and metaphase, which we term the time of spindle assembly, 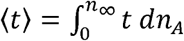, where *n*_∞_= lim_*t* → ∞_ *n*_A_(t).

On a single cell level, we model the relevant processes of spindle assembly. We calculate the rate of KC capture by taking into account known microtubule dynamics and geometry of yeast spindles (Figure 1B). Microtubules nucleate from the spindle pole body at rate *v*_i_ and extend towards the spindle equator. They can attach to an unattached KC with probability *P*. The rate of KC attachment is the probability of attachment of one of the unattached KCs multiplied with the microtubule nucleation rate, and for *C* – i unattached KCs reads

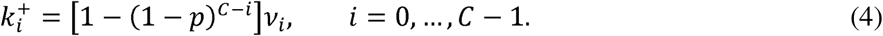

For other values of the index *i* the rate of KC attachment is zero to exclude unrealistic cases, with negative number of chromosomes or with more than *c* chromosomes. We calculate the nucleation rate at the spindle pole body as *v*_i_ =*v* · (*M* – *i*), where we take into account that the spindle pole body with *M* nucleation sites has *M* – *i* unoccupied nucleation sites and the nucleation rate for one nucleation site is *v*. In our model, attachment occurs when a microtubule contacts the KC. The probability of attachment is calculated based on spindle geometry as the ratio of the cross-section areas of the KC, *s*_KC_, and the total area of the spindle, *p*= *s*_KC_/(*SM* + *s*_KC_). Here *s* denotes the cross-section area occupied by one microtubule. To consider cases with different numbers of chromosomes, we introduce a linear relationship between the number of chromosomes and nucleation sites, *M*= α*c* + 4, which is based on experimental findings (Nannas et al., 2014; Storchova et al., 2006). The parameter α is typically around 1. Microtubules detach from one KC at constant rate *k*- and in the case of *i* attached KCs, the detachment rate is given by

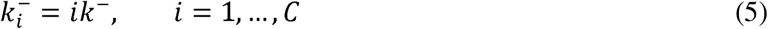

whereas the detachment for other values of the index *i* is zero. Cells proceed from metaphase to anaphase by silencing the spindle assembly checkpoint, but they can also proceed from prometaphase to anaphase when they spend a prolonged time in mitotic arrest, which in our model results in chromosome loss (Rieder et al., 1994; Rudner and Murray, 1996). We distinguish these two cases by introducing a rate of anaphase entry given by

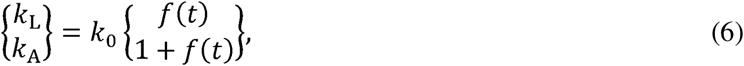

where in the upper and bottom rows we calculate rates for cells in prometaphase and metaphase, respectively. We describe bypassing the checkpoint in mitotic arrest with a function of time *f(t)*, irrespective whether cells are in prometaphase of metaphase. Because this function has not been studied so far, we choose a simple mathematical form *f(t) = exp*[(*t-t*_0_*)/t*_0_], which takes into account that rate of anaphase entry increases in time. Here, parameters *t*_0_ and *t*_c_ denote the duration of mitotic arrest and the variability of this process, respectively. Cells can also enter anaphase by silencing the checkpoint, which happens for the fraction of cells in metaphase at a constant rate, *k*_0_.

## SUPLEMENTAL INFORMATION

### SUPPLEMENTAL FIGURES

**Figure S1.**
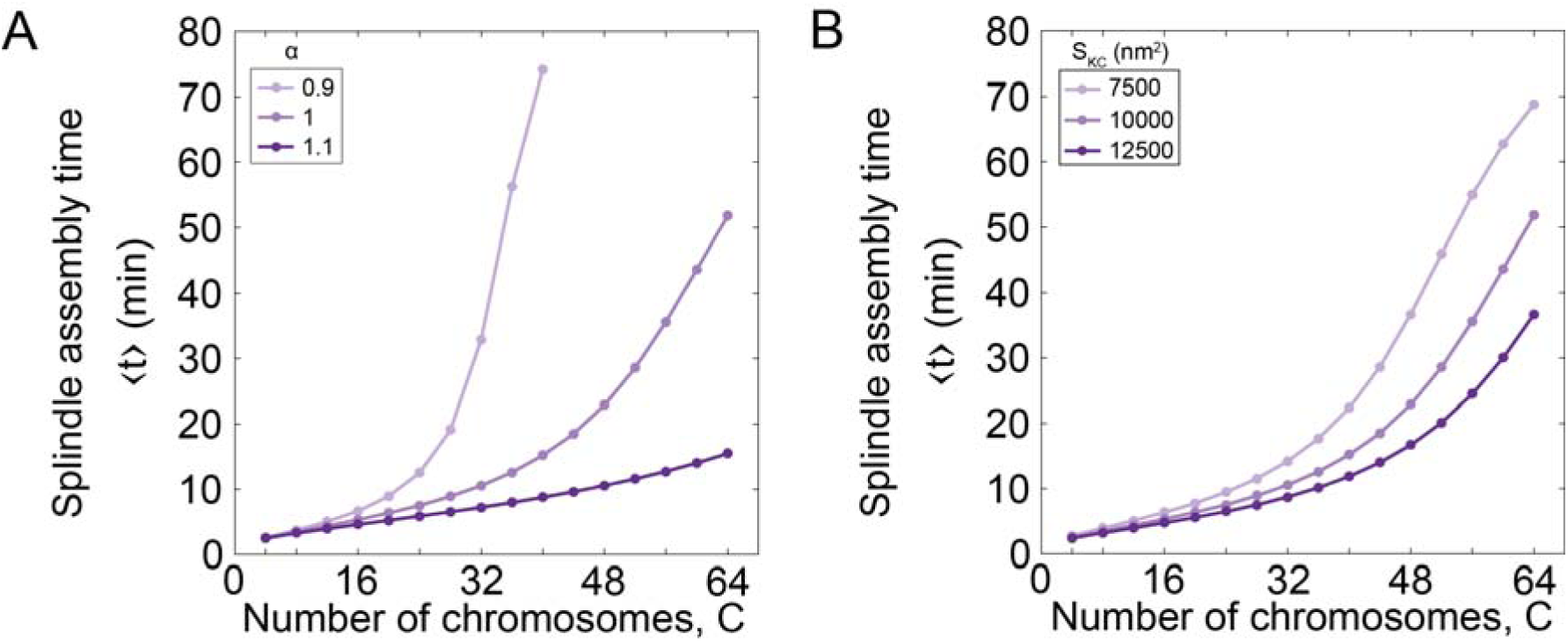
Related to Figure 2C. Time of spindle assembly for different number of chromosomes and different values of model parameters. (A) As our model describes spindle formation, we explore the relevance of parameters on the time of spindle assembly. We varied the number of chromosomes and the parameter that links the number of chromosomes and microtubule nucleation sites, *α*. We found that the time of spindle assembly increases with the number of chromosomes. For parameter values *α= 1.0* the time of spindle assembly increases with the number of chromosomes. By increasing *α* to values greater than 1 the assembly speeds up for a larger number of chromosomes. By decreasing the parameter to the value *α = 0.9* the assembly time dramatically increases with number of chromosomes and goes to infinity when there are more than 40 chromosomes. The infinite time of spindle assembly occurs for cells in which the number of microtubule nucleation sites at one pole is smaller than number of chromosomes. Interestingly, in yeast the value of the parameter *α* in cells is close to 1 (Table 1). Three different shades correspond to different number of chromosomes, *α* =0.9, 1.0, 1.1. For color-codes see inset legend. The other parameters are given in Figure 1C. (B) We also explored the relevance of geometry by varying the cross-section area of the KC, *s*_KC_. We found that geometry has a small contribution for small number of chromosomes, but for larger number of chromosomes, the time of spindle assembly decreases with the cross-section area. The role of the cross-section area occupied by one microtubule can be inferred from these data because both parameters contribute to attachment probability *P*. Three different shades correspond to different cross-section area of the KC, *s*_KC_ = 7500 nm^2^, 10000 nm^2^, 12500 nm^2^. For color-codes see inset legend. The other parameters are given in Figure 1C.

**Figure S2.**
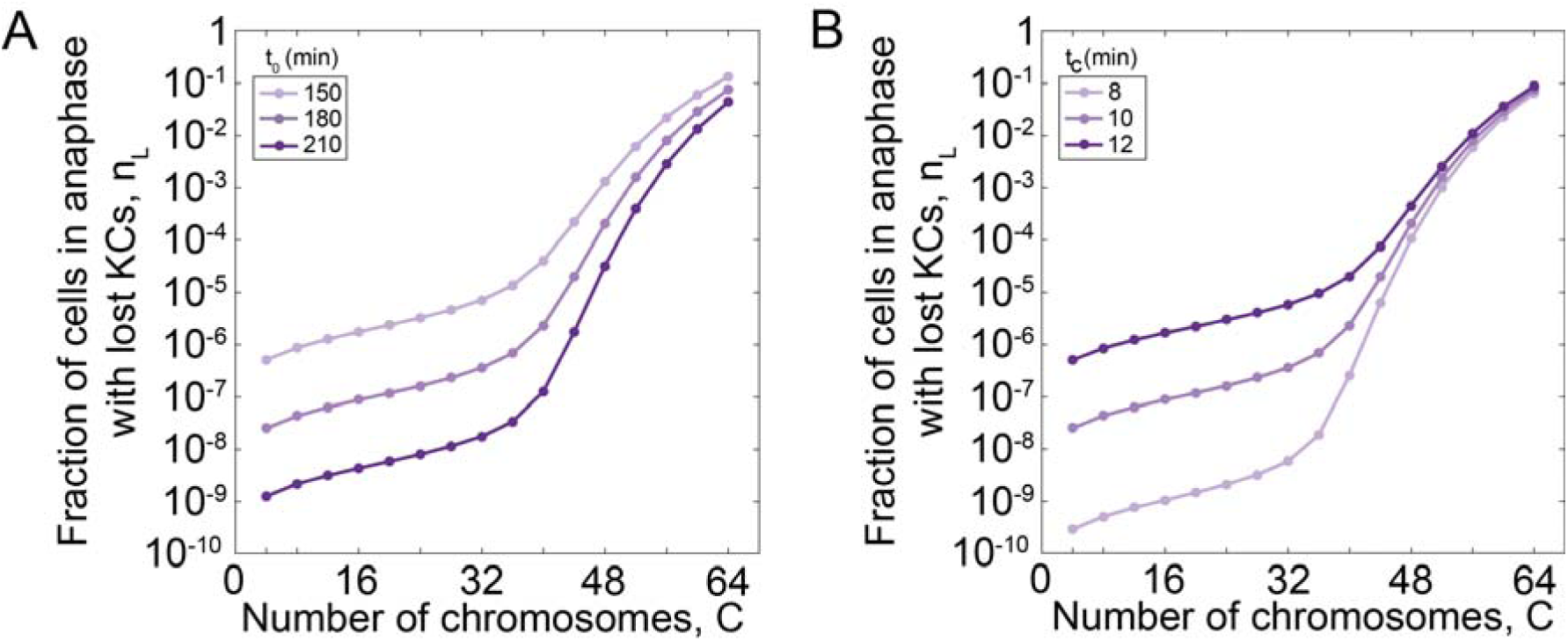
Related to Figure 2D. Rate of chromosome loss for different number of chromosomes and different values of parameters describing the maximum duration of mitotic arrest used in function *f*(*t*). A) Rate of chromosome loss for different values of parameter that describe the duration of mitotic arrest, *t*_0_. We find that cells with shorter duration of mitotic arrest have an increased chromosome loss rate, which increases for cells with higher ploidy, such as tetraploid cells, irrespective of parameter *t*_0_. Three different shades correspond to different values of parameter *t*_0_ = 150 min, 180 min,210 min. For color-codes see inset legend. The other parameters are given in Figure 1C. (B). Rate of chromosome loss for different values of parameter that describe variability in duration of mitotic arrest, *t*_0_. We find that cells with a smaller variability in mitotic arrest have a smaller rate of chromosome loss. This trend remains for cells with different number of chromosomes. Three different shades correspond to different values of parameter *t*_0_ = 8 min,10 min,12 min. For color-codes see inset legend. The other parameters are given in Figure 1C.

## REFERENCES

Akiyoshi, B., Sarangapani, K.K., Powers, A.F., Nelson, C.R., Reichow, S.L., Arellano-Santoyo, H., Gonen, T., Ranish, J.A., Asbury, C.L., and Biggins, S. (2010). Tension directly stabilizes reconstituted kinetochore-microtubule attachments. Nature 468, 576–579.

Comai, L. (2005). The advantages and disadvantages of being polyploid. Nat Rev Genet 6, 836–846.

Gardner, M.K., Bouck, D.C., Paliulis, L.V., Meehl, J.B., O’Toole, E.T., Haase, J., Soubry, A., Joglekar, A.P., Winey, M., Salmon, E.D., et al. (2008). Chromosome congression by Kinesin-5 motor-mediated disassembly of longer kinetochore microtubules. Cell 135, 894–906.

Gay, G., Courtheoux, T., Reyes, C., Tournier, S., and Gachet, Y. (2012). A stochastic model of kinetochore-microtubule attachment accurately describes fission yeast chromosome segregation. J Cell Biol 196, 757–774.

Gerstein, A.C., Chun, H.J., Grant, A., and Otto, S.P. (2006). Genomic convergence toward diploidy in Saccharomyces cerevisiae. PLoS Genet 2, e145.

Gonen, S., Akiyoshi, B., Iadanza, M.G., Shi, D., Duggan, N., Biggins, S., and Gonen, T. (2012). The structure of purified kinetochores reveals multiple microtubule-attachment sites. Nat Struct Mol Biol 19, 925–929.

Hill, T.L. (1985). Theoretical problems related to the attachment of microtubules to kinetochores. Proc Natl Acad Sci U S A 82, 4404–4408.

Hufton, A.L., and Panopoulou, G. (2009). Polyploidy and genome restructuring: a variety of outcomes. Curr Opin Genet Dev 19, 600–606.

Kalinina, I., Nandi, A., Delivani, P., Chacon, M.R., Klemm, A.H., Ramunno-Johnson, D., Krull, A., Lindner, B., Pavin, N., and Tolic-Norrelykke, I.M. (2013). Pivoting of microtubules around the spindle pole accelerates kinetochore capture. Nat Cell Biol 15, 82–87.

Li, R., and Murray, A.W. (1991). Feedback control of mitosis in budding yeast. Cell 66, 519–531.

Lu, Y.J., Swamy, K.B., and Leu, J.Y. (2016). Experimental Evolution Reveals Interplay between Sch9 and Polyploid Stability in Yeast. PLoS Genet 12, e1006409.

Mayer, V.W., and Aguilera, A. (1990). High levels of chromosome instability in polyploids of Saccharomyces cerevisiae. Mutat Res 231, 177–186.

Mitchison, T.J., and Kirschner, M.W. (1985). Properties of the kinetochore in vitro. II. Microtubule capture and ATP-dependent translocation. J Cell Biol 101, 766–777.

Nannas, N.J., O’Toole, E.T., Winey, M., and Murray, A.W. (2014). Chromosomal attachments set length and microtubule number in the Saccharomyces cerevisiae mitotic spindle. Mol Biol Cell 25, 4034–4048.

O’Toole, E.T., Mastronarde, D.N., Giddings, T.H., Jr., Winey, M., Burke, D.J., and McIntosh, J.R. (1997). Three-dimensional analysis and ultrastructural design of mitotic spindles from the cdc20 mutant of Saccharomyces cerevisiae. Mol Biol Cell 8, 1–11.

Otto, S.P., and Whitton, J. (2000). Polyploid incidence and evolution. Annu Rev Genet 34, 401–437.

Paul, R., Wollman, R., Silkworth, W.T., Nardi, I.K., Cimini, D., and Mogilner, A. (2009). Computer simulations predict that chromosome movements and rotations accelerate mitotic spindle assembly without compromising accuracy. Proc Natl Acad Sci U S A 106, 15708–15713.

Rieder, C.L., Schultz, A., Cole, R., and Sluder, G. (1994). Anaphase onset in vertebrate somatic cells is controlled by a checkpoint that monitors sister kinetochore attachment to the spindle. J Cell Biol 127, 1301–1310.

Rudner, A.D., and Murray, A.W. (1996). The spindle assembly checkpoint. Curr Opin Cell Biol 8, 773–780.

Schulman, I.G., and Bloom, K. (1993). Genetic dissection of centromere function. Mol Cell Biol 13, 3156–3166.

Selmecki, A.M., Maruvka, Y.E., Richmond, P.A., Guillet, M., Shoresh, N., Sorenson, A.L., De, S., Kishony, R., Michor, F., Dowell, R., et al. (2015). Polyploidy can drive rapid adaptation in yeast. Nature 519, 349–352.

Storchova, Z., Breneman, A., Cande, J., Dunn, J., Burbank, K., O’Toole, E., and Pellman, D. (2006). Genome-wide genetic analysis of polyploidy in yeast. Nature 443, 541–547.

Tubman, E.S., Biggins, S., and Odde, D.J. (2017). Stochastic Modeling Yields a Mechanistic Framework for Spindle Attachment Error Correction in Budding Yeast Mitosis. Cell Syst 4, 645–650 e645.

Vasileva, V., Gierlinski, M., Yue, Z., O’Reilly, N., Kitamura, E., and Tanaka, T.U. (2017). Molecular mechanisms facilitating the initial kinetochore encounter with spindle microtubules. J Cell Biol 216, 1609–1622.

Volkov, V.A., Zaytsev, A.V., Gudimchuk, N., Grissom, P.M., Gintsburg, A.L., Ataullakhanov, F.I., McIntosh, J.R., and Grishchuk, E.L. (2013). Long tethers provide high-force coupling of the Dam1 ring to shortening microtubules. Proc Natl Acad Sci U S A 110, 7708–7713.

Winey, M., Mamay, C.L., O’Toole, E.T., Mastronarde, D.N., Giddings, T.H., Jr., McDonald, K.L., and McIntosh, J.R. (1995). Three-dimensional ultrastructural analysis of the Saccharomyces cerevisiae mitotic spindle. J Cell Biol 129, 1601–1615.

Wollman, R., Cytrynbaum, E.N., Jones, J.T., Meyer, T., Scholey, J.M., and Mogilner, A. (2005). Efficient chromosome capture requires a bias in the ‘search-and-capture’ process during mitotic-spindle assembly. Curr Biol 15, 828–832.

Zaytsev, A.V., and Grishchuk, E.L. (2015). Basic mechanism for biorientation of mitotic chromosomes is provided by the kinetochore geometry and indiscriminate turnover of kinetochore microtubules. Mol Biol Cell 26, 3985–3998.

